# Hyperpolarization-activated cation channels confer tonotopic specialization for temporal encoding of sound frequency in the cochlear nucleus

**DOI:** 10.1101/2025.06.03.657729

**Authors:** Kwame Owusu-Nyantakyi, Lashaka S. Hamlette, Go Ashida, Sonia Weimann, Stefan N. Oline, R. Michael Burger

**Affiliations:** Department of Biological Sciences, Lehigh University, Bethlehem, Pennsylvania 18015; Cluster of Excellence "Hearing4all," Department of Neuroscience, University of Oldenburg, 26129 Oldenburg, Germany

## Abstract

Sensory neurons are equipped with physiological properties vital for accurate signal processing. The functional importance of such properties is exemplified in auditory circuits where intrinsic excitability is optimized to detect frequency-specific features. In birds, the neurons of nucleus magnocellularis (NM) receive primary auditory input (Rubel and Parks, 1975a; Parks and Rubel, 1978; Jackson et al., 1982) and are arranged tonotopically. NM comprises a superficially homogenous neural population, but several physiological properties vary systematically along its tonotopic frequency axis. In particular, expression of voltage-gated conductances plays a pivotal role in creating selectivity that enables temporal precision. Here, we identify a previously undescribed gradient of hyperpolarization-activated cation channels (IH). Whole cell patch clamp techniques and immunostaining for HCN1, an IH channel subunit, demonstrated an expression gradient corresponding to NM’s tonotopic axis. To investigate the function of tonotopic IH expression in NM, we applied a depolarizing ramp injection protocol to measure the impact of pharmacologically blocking IH on neural active properties (Ferragamo and Oertel, 2002; McGinley and Oertel, 2006; Oline et al. 2016). Next, we investigated whether this tonotopic patterning of HCN facilitates encoding of temporally patterned inputs. We injected depolarizing current pulse trains before and during HCN channel block. During pharmacological block, there was a reduction of NM spike entrainment to input pulses suggesting a key contribution of HCN channels to NM’s ability to encode its synaptic drive. Results show that there is tonotopic distribution of HCN channels in NM which provides a novel mechanism that enables NM neurons to encode temporally patterned excitatory input.

**Significance Statement:** This study is the first to describe a tonotopic gradient of IH channels in a vertebrate cochlear nucleus. Physiological and computational model assays suggest that the tonotopic expression pattern of HCN channels enables improved neural encoding of high frequency, temporally patterned input. Temporal response fidelity enables precise sound localization computations.

## Introduction

In vertebrates, neural centers devoted to processing sensory information are often organized into topographic maps associated with specific stimulus parameters. Auditory regions exhibit tonotopy, the topographic representation of sound frequency which recapitulates the organization of the auditory sensory epithelium. In the cochlea, tonotopically arranged hair cells are innervated by bipolar auditory nerve fibers, which project to the brainstem cochlear nucleus. In birds, nucleus magnocellular is (NM) preserves and sharpens temporal information present in auditory nerve responses that is essential for sound location computations (Parks and Rubel, 1978; Lippe and Rubel, 1985; Fukui et al., 2006). Temporal information is encoded by phase-locked action potentials that reliably discharge at a particular phase of the stimulus waveform. High frequency stimuli challenge phase-locking in two ways. First, spiking driven by high frequency sound skips stimulus cycles as neurons near their maximum firing rate. Second, the short duration of each stimulus period for high frequencies falls below neuronal capacity to sustain temporal precision. Despite these constraints, NM neurons achieve significant phase-locked precision at frequencies exceeding 2 kHz, despite failure to entrain responses to every stimulus cycle (Joris et al., 1994; Fukui and Ohmori, 2004). This remarkable computational ability is rooted in several tonotopically distributed specializations including patterned expression of voltage-gated ion channels (Fukui and Ohmori, 2004). NM expresses gradients of both low and high voltage gated potassium channels, KLVA and KHVA, respectively. These channels tune responses in the frequency domain by imparting differential filtering properties along the tonotopic axis (Oline et al., 2016; Hong et al., 2018). KLVA is preferentially expressed in high characteristic frequency (HCF) neurons, and improves phase-locking by shunting poorly-timed inputs (Kopp-Scheinpflug, 2017; Lu et al., 2017), while KHVA expression reduces spike half-width and supports firing at very high rates (Parameshwaran-Iyer et al., 2003; Lu et al., 2004; Lu et al., 2017). Increased potassium conductances also endow HCF neurons with low input resistance, which facilitates phase-locking by shunting poorly timed synaptic input (Monsivais et al., 2000). Low characteristic frequency (LCF) neurons have relatively higher input resistance which enables temporal precision to benefit from integration of smaller, subthreshold inputs (Oline et al., 2016).

The high potassium channel expression in NM lowers resting potentials into the activation range of a second class of voltage-gated cation channels, the hyperpolarization-activated cyclic-nucleotide gated cation current (*I*H) channels (Ramakrishnan et al., 2012). Although *I*H channels exhibit slow kinetics and are therefore unlikely to modulate auditory responses on a cycle-by-cycle basis, they have been shown to contribute to temporal processing in auditory neurons (Bal and Oertel, 2001; Fischl et al., 2016). *I*H conductances are partially activated near rest, with a reversal potential near ∼-35 mV (Santoro and Tibbs,1999). Depolarization by IH current partially offsets hyperpolarization by KLVA channels. Co-expression of these channels maintains rest near spike threshold by lowering the input resistance. This facilitates synaptic integration which is crucial for temporal precision (Yamada et al., 2005; Lewis et al., 2010). In binaural "coincidence detecting" medial superior olive neurons (MSO) or nucleus laminaris (NL) in mammals and birds, tonotopic gradients of IH have been shown to contribute to temporal precision (Yamada et al., 2005; Baumann et al., 2013). Complementary deactivation kinetics of *I*H and *I*K-LVA in the MSO are balanced such that responses to depressing synaptic drive are maintained by a steady increase in total membrane input resistance (Khurana et al., 2011). Finally, the stochastic gating of *I*H contributes significantly to membrane voltage noise which has been shown to positively influence spike timing reliability in cortical regions (Schneidman and Segev, 1998; White et al., 2000; Kole et al., 2006).

Here we report physiological, immunohistochemical, and computational modeling studies investigating the distribution and function of *I*H conductances in NM neurons across the tonotopic axis. We show that temporally precise responses to simulated synaptic input across NM is *I*H -dependent, especially at high input rates. Correspondingly, we show that *I*H channel expression is most prominent in the HCF region. Modeling further suggests that *I*H reduces error responses in HCF neurons induced by stochastic membrane fluctuations. Together these results suggest that *I*H plays an essential role in conditioning membrane excitability to optimize temporal encoding of auditory stimuli across frequencies.

## Materials and Methods

### Subjects

Fertilized chicks (*Gallus gallus domesticus*) of either sex were obtained from a local hatchery (Hoover’s Hatchery) and were raised at Lehigh University’s central animal facility. All protocols were approved by the Lehigh University Institutional Animal Care and Use Committee and were performed in compliance with the Public Health Service Policy on Humane Care and Use of Laboratory Animals. We recorded from NM neurons at age E16-E19,

### Slice preparation

Brain slices were prepared as described in Oline and Burger (2014). Briefly, embryos aged E15-E19 (n = 39) were removed from eggs followed by a rapid decapitation. Hatched chicks were anesthetized using isoflurane (MINRAD). 150-microncoronal brain slices were collected using a vibrating microtome (Campden Instruments 7000 smz2), and slices were stored in oxygenated artificial cerebral spinal fluid (aCSF) containing (in mM): 130 NaCl, 3 KCl, 10 glucose, 1.25 NaH2PO4, 26 NaHCO3, 3 CaCl2, and 1 MgCl2 at 22C. Slices were kept in order in a slice chamber with constantly oxygenated aCSF and incubated for 45 minutes at 37°C. For each brain, 7-8 slices in total were collected. The tonotopic position was estimated according to position along the known rostromedial (high CF) to caudolateral (low CF) axis (Rubel and Parks, 1975; Fukui and Ohmori, 2004). We utilized the tonotopic characterization scheme established by Fukui and Ohmori (2004) that divides NM into 11 tonotopically defined regions from LCF to HCF. Thus, for our physiology experiments, low CF cells from the first slice and the lateral third of the second slice, and high CF cells from in the last slice and the medial third of the second-to-last slice were used.

### Recording arrangement

Electrophysiological experiments were made using a custom recording chamber on a retractable chamber shuttle system (Siskiyou Corporation), and neurons were visualized with a Nikon FN-1 Physiostation microscope. The recording chamber was continuously superfused with oxygenated aCSF. Slice temperature was monitored and maintained at 35 ± 0.50°C using an inline feedback temperature controller and heated stage (Warner Instruments). NM neurons are easily identifiable within the clear boarders of NM. Whole-cell voltage-clamp and current-clamp recordings were made using an Axon Instruments Multiclamp 700B amplifier. The signals were digitized with a Digidata 1440 data acquisition board and recorded using the Clampex software by Molecular Devices. Patch pipettes were pulled using a two-stage Narishige puller (model PC-10) with thick-walled borosilicate glass capillary tubes (1B120F-4, World Precision Instruments). Pipettes were filled with a potassium-based solution containing (in mM): 145 K-gluconate, 5 KCl, 1 MgCl2, 10 HEPES, and 5 EGTA, pH adjusted to 7.2 with KOH. Junction potential was empirically determined at 10 mV, data are presented uncorrected. For both current and voltage-clamp recording, a test solution was applied to the bath to block GABA and AMPA receptors (500 nM SR95531, and 40 µm DNQX) (Tocris Bioscience). To block *I*H, 40 µM ZD-7288 (Tocris Bioscience) was bath applied.

### Code Accessibility

Custom code was created using MATLAB to analyze current-clamp and voltage-clamp data and are readily available upon request to the authors.

### Data Analysis

For pulse train experiments, we set pulse amplitude to each neuron’s rheobase value. Rheobase was defined as the minimum amplitude depolarizing square current (50 ms) required to evoke a spike in at least 50% of trials. Membrane excitability was assessed using slope threshold, integration period and action potential halfwidth for the lowest threshold spike in response to systematically varying ramp stimuli (Oline et al., 2016). Slope threshold was defined as the slope of the voltage response (mV/ms) during the middle 20%-80% amplitude range between ramp onset and spike threshold (Ferragamo and Oertel, 2002; McGinley and Oertel, 2006). Integration period was defined as the latency from ramp onset to spike initiation. Action potential halfwidth was defined as action potential duration in ms at 50% of peak amplitude. To analyze the magnitude of the depolarizing voltage sags observed in response to hyperpolarization, we calculated the difference between the maximum negative voltage deflection evoked by the largest hyperpolarizing current injection and the average membrane voltage during the last 10 ms of the hyperpolarizing step. The sag magnitude difference was then expressed as a percentage (**Fig. 1 C, F)**. To analyze the *I*H current magnitude across NM, we subtracted control traces from traces recorded following bath application of ZD-7288. To quantify the slowly gating *I*H magnitude, we subtracted the instantaneous voltage step onset-amplitude from the steady-state current at each holding voltage during long duration (800 ms) voltage steps. We observed that cell membrane integrity was challenged by hyperpolarization below -120 mV. Therefore, our protocols limited hyperpolarizing steps to -120 mV. A one-way repeated measures ANOVA was conducted for the current amplitudes produced during the activation range of *I*H between low and high CF neurons. A paired t-test was used to determine differences in the integration period and slope thresholds for each group. We injected rectangular current pulses at 100 and 200Hz for 2 seconds before and after blocking *I*H and measured the entrainment response in the last 1.9 s of stimulus train input.

**Figure 1.**
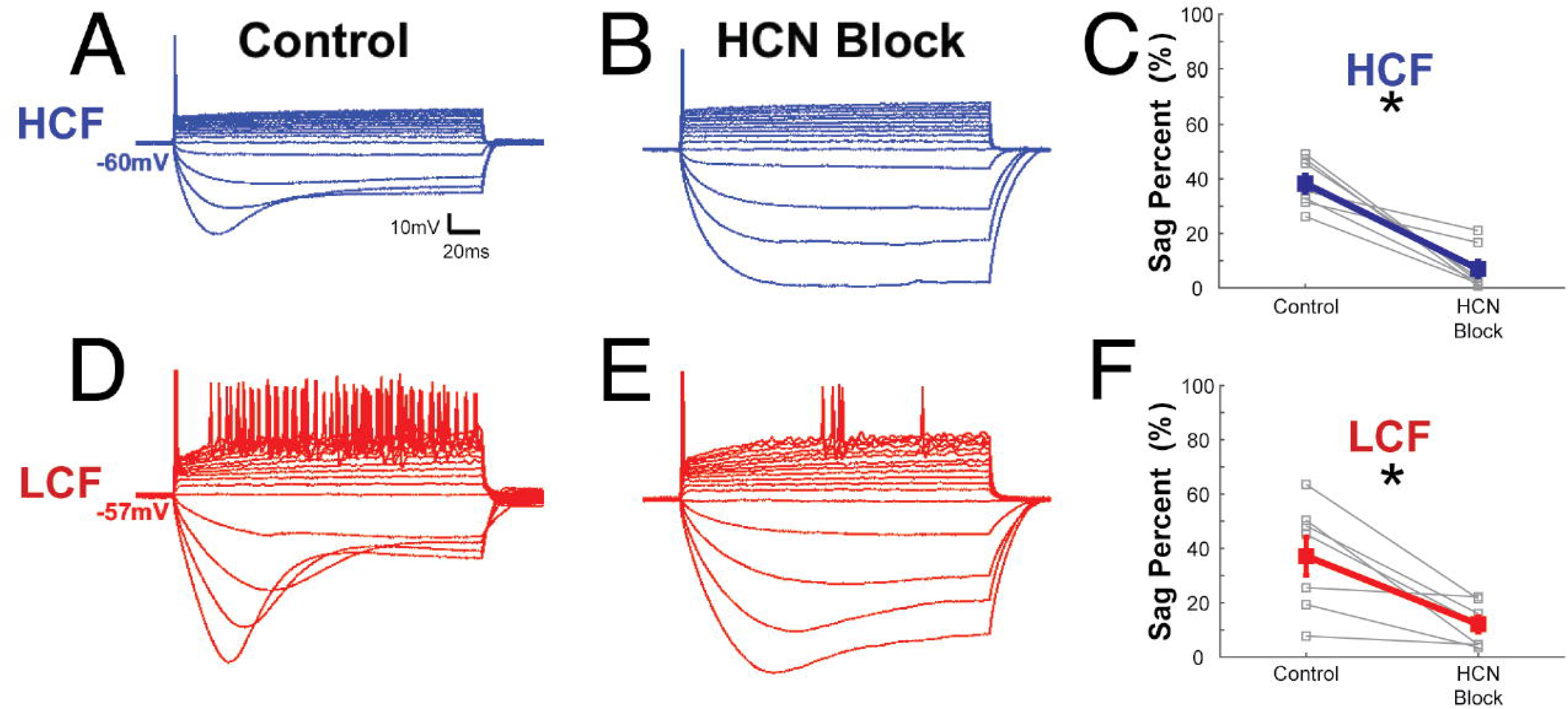
Hyperpolarizing current injection to NM neurons induces a depolarizing voltage sag that is IH-dependent. A) Membrane voltage recordings of HCF, and D) LCF NM neurons to current injections in control treatment. B) Voltage responses of the same HCF and E) LCF NM neurons during ZD7288 treatment. C&F) Sag percent change in control and during application of ZD7288 for HCF and LCF neurons.

### Immunohistochemistry

Twelve chickens age P3 were used for immunohistochemical (IHC) methods, when most auditory circuitry is structurally and functionally mature (Rubel and Fritzsch, 2002). Chicks were deeply anesthetized with an overdose of Somnasol veterinary euthanasia solution (Henry Schein Animal Health: >0.25 ml/kg body weight) and transcardially perfused with PBS followed by 4% paraformaldehyde (PFA) in PBS, pH 7.4. Brains were removed and postfixed in 4% PFA overnight at 4°C then rinsed, the auditory brainstem was blocked. To create slices containing the full tonotopic range of NM, we sectioned tissue at 50 µm parallel to the main tonotopic caudolateral to rostromedial extent of NM. (Parks and Rubel, 1975). (7000 smz2, Camden Instruments). Tissue was rinsed, blocked in 10% normal goat serum for 1h, then incubated overnight at 4°C in solution containing 5% normal goat serum, 0.1 % triton-x, and anti-HCN1 (1:200; Alomone labs, cat. # APC-056). Sections were rinsed and then incubated for 2 hours with secondary antibodies (1:300; ThermoFisher Scientific) conjugated to Alexa Fluor 488 goat anti-rabbit to label. Sections were mounted on slides and counterstained with nissl solution (1:200; Invitrogen) then mounted in Vectashield containing DAPI medium, and confocal images were captured (LSM 880 Meta, Zeiss). No staining was observed when primary antibodies were absent (*data not shown*) or when HCN1 antibody was preabsorbed with antigen peptide (**Fig. 3D, I**). Pixel intensity of NM neurons was quantitatively analyzed using Zeiss software (Zen lite 3.11) by creating regions of interest around individual neurons. Pixel intensity was normalized to the highest value in each slice and data was averaged across ten neurons from each tonotopic region in four different animals. Significance was measured using a paired t-test. Adobe Photoshop and Illustrator Software were used to label, colorize, and crop images for display.

**Figure 2.**
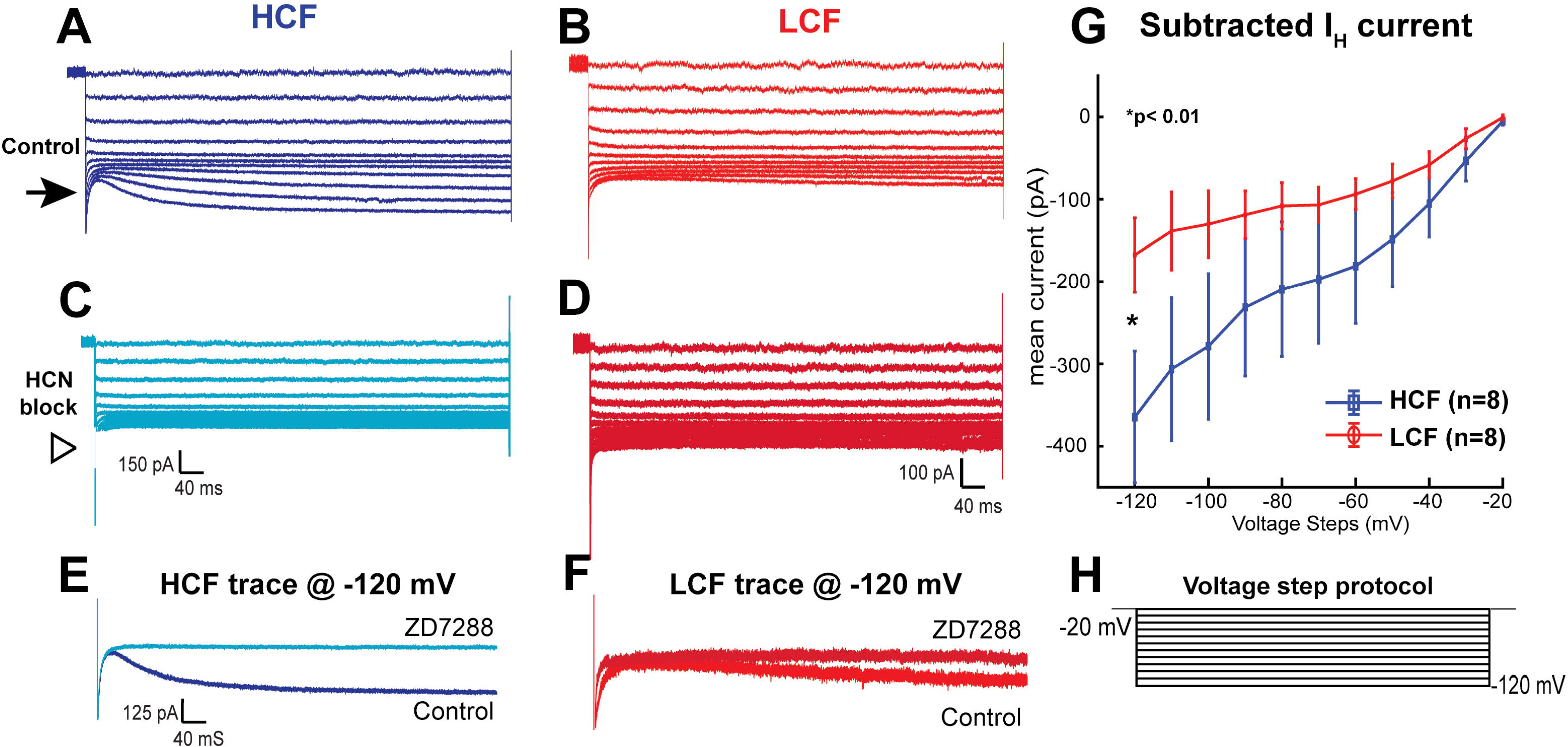
HCN conductances are enriched in HCF compared to LCF neurons. A&B) Voltage clamp responses of HCF and LCF NM neurons to long duration voltage steps (−20 to −120 mV) before ZD7288 application, and *C&D*) during ZD7288 treatment. In both neurons a slowly-developing inward current, indicative of HCN conductance, is exhibited at hyperpolarizing voltage steps (*black arrows in A&B*) which are reduced in the presence of ZD7228 treatment (*open arrowheads* in *C&D*). E&F) show current responses of the same neurons at the maximum negative holding voltage (−120 mV) before and during ZD7288 application indicating a relatively stronger conductance in the HCF neuron. G) shows an I-V plot showing the population means for HCN dependent current responses of both cell types at each holding voltage. Data confirm larger conductances in HCF neurons over the entirety of the HCN channel voltage activation range. *p=0.0031 (two-way ANOVA). H) command voltage step protocol.

**Figure 3.**
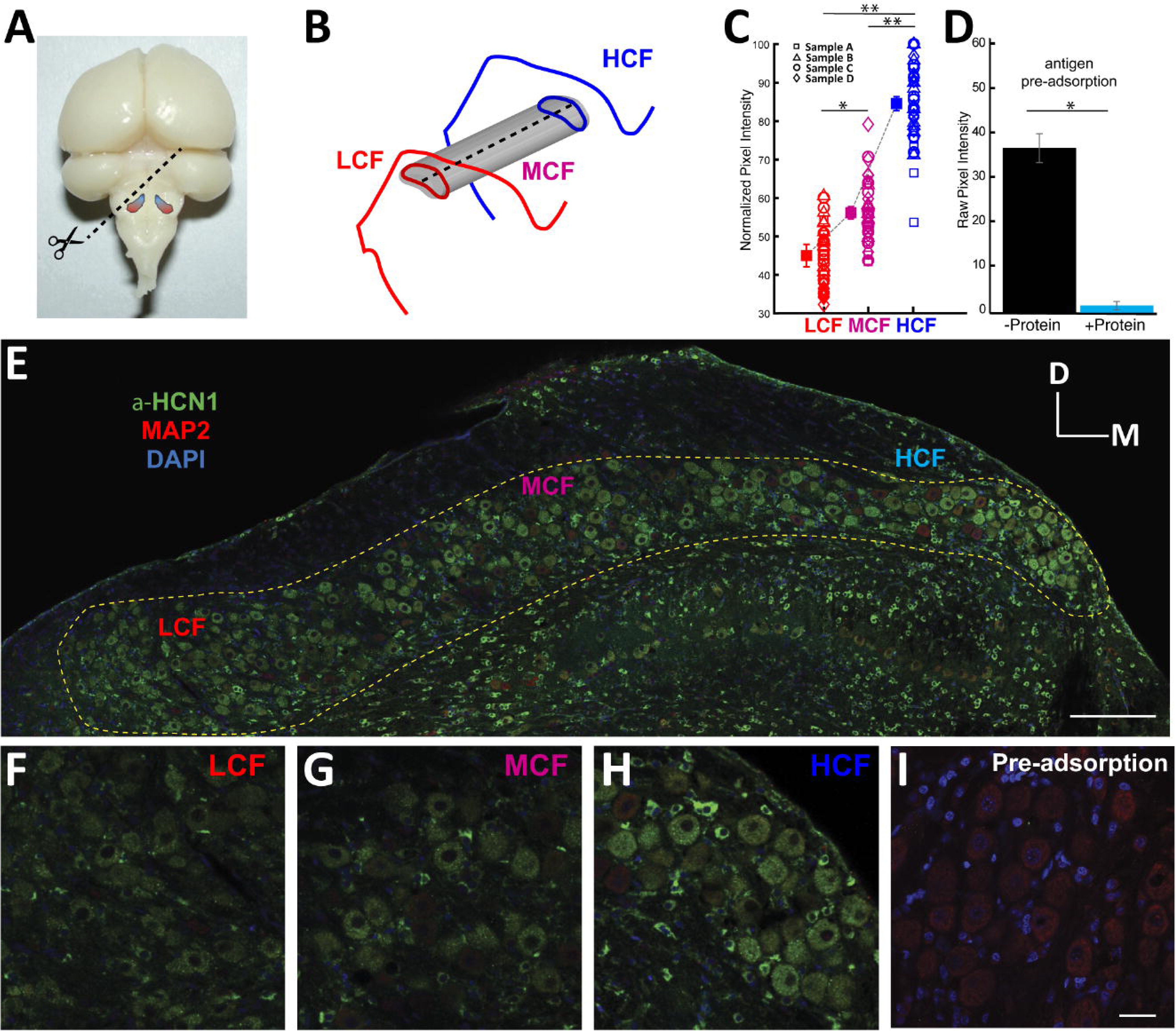
HCN1 channels exhibit stronger expression in high frequency regions of NM. A) Dorsal view of tonotopic organization of NM in chick brain (*blue* indicates high frequency and *red* indicates low frequency). The dotted line indicates para-tonotopic slice orientation. B) Illustration of tonotopic regions in NM. *Red* indicates LCF, *magenta* indicates MCF, and *blue* indicates HCF. C) Pixel intensity values quantitatively indicate that HCN1 expression is stronger in HCF regions of NM when compared to MCF and LCF. Regions of interest were created around 10 individual NM neurons from each frequency region in four different samples (ο = sample A, △ = sample B, ϒ = sample C, ↓ = sample D). D) Pixel intensity values indicate that protein antigen pre-adsorption nearly abolished HCN1 staining (*blue bar*) when compared to control (*black bar*). E) Tiled image depicting immunofluorescence of HCN1 staining in a 50 µm para-tonotopic slice of NM. *Dashed line* indicates NM region in the slice. *Green* indicates anti-HCN1 stain, *red* indicates anti-MAP2 stain, and *blue* indicates DAPI stain. F) LCF, and G) MCF demonstrate lower HCN1 staining compared to H) HCF. Immunostaining for HCN1 was absent in tissue processed without primary antibody (data not shown) I) Illustration of pre-adsorbed antibody demonstrating no anti-HCN1 staining in P3 tissue. Images were obtained with confocal microscopy at 40X magnification. Scale bars, 200 µm (E) and 50 µm (I).

### NM modeling

Using voltage clamp recordings from chick NM neurons, we developed a computational model of low and high frequency neuronal subtypes that include IH current. (**Table 1; Supplementary** Fig. 1A). The equations and parameters of the model are summarized in Tables 1 and 2, respectively. The HCF model has a larger *I*H conductance than the LCF model (Supplementary Fig. 1B; compare it with experimental **Fig. 5E**), while the activation kinetics of the *I*H current is identical between these two models. We combined this *I*H model with our previous NM model (Oline et al., 2016) to investigate possible roles for the *I*H current. First, we tested our LCF and HCF models by applying the same ramp currents as in our slice experiments (**Fig. 1C, D**). Application of the *I*H channel blocker was simulated by a removal of *I*H current (i.e., zero conductance). As in slice recordings **(experimental Fig. 2C, D**), the LCF model consistently shows a lower slope threshold **(Supplementary** Fig. 1E) and a longer integration window **(Supplementary** Fig. 1F) at each holding potential than the HCF model. However, the simulated differences between the control and ZD conditions were very small. We regarded these marginal differences as acceptable, considering the larger neuron-to-neuron variability than the actual effect size **(experimental Fig. 2C, D**).

**Figure 4.**
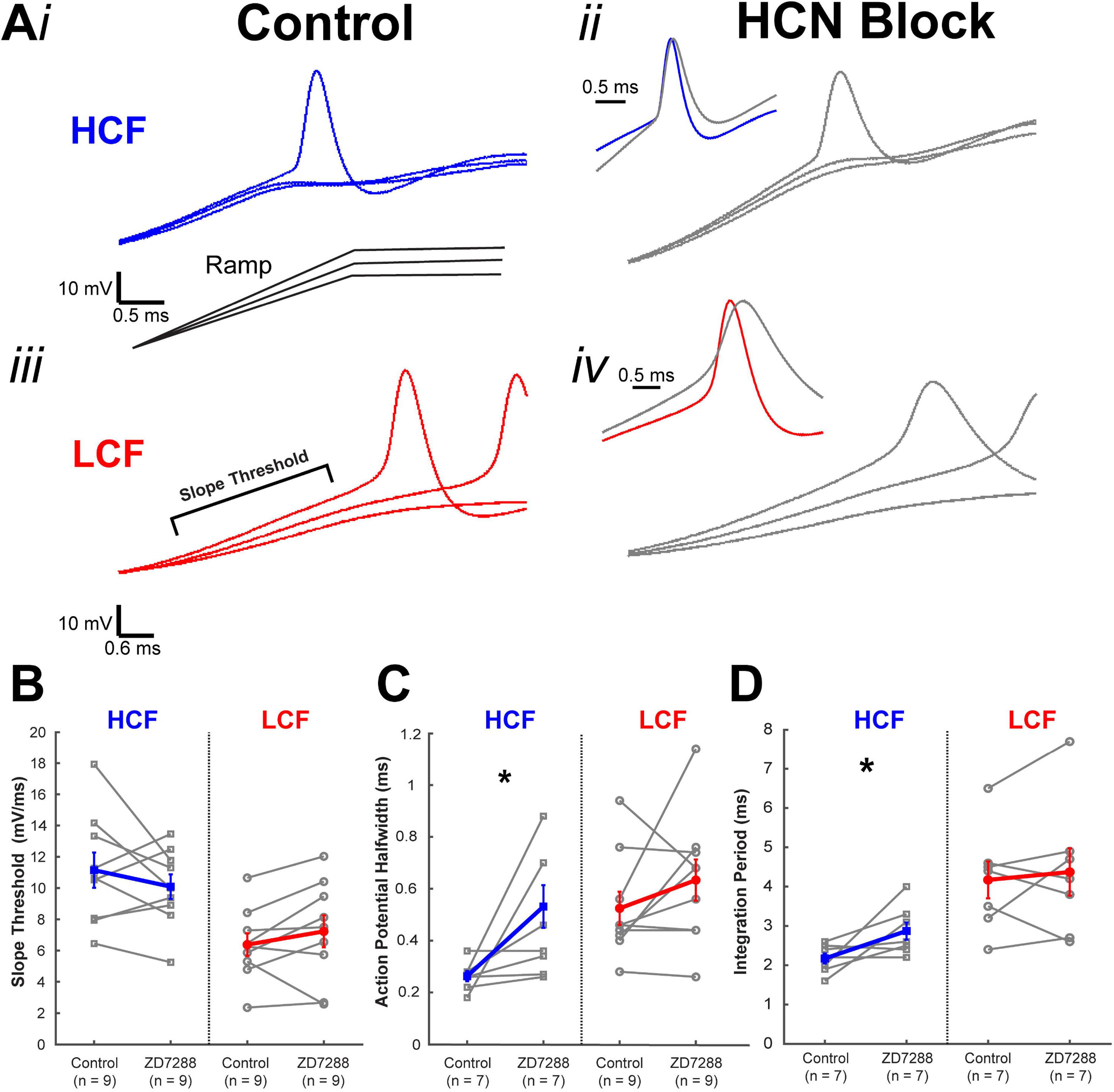
HCN channels modulate integration period and action potential halfwidth in HCF neurons. A*i*) Representative membrane voltage response of an HCF and A*iii*) LCF neuron to ramp current injection before ZD7288 application. A*ii*) Membrane voltage response to ramp current injection of HCF and, A*iv*) LCF neurons during ZD7288 application. *Insets* show the pre and post drug traces for the lowest threshold spikes where increased halfwidth is observable in post drug *grey* traces. B) Slope threshold did not change for either population in the presence of ZD7288. C) Action potential halfwidth was significantly greater for HCF neurons (*p=0.0122, Students t-test), but not for LCF neurons. D) Similarly, integration period was longer for HCF neurons (*p=0.0255, Students t-test) but not for LCF neurons.

**Figure 5.**
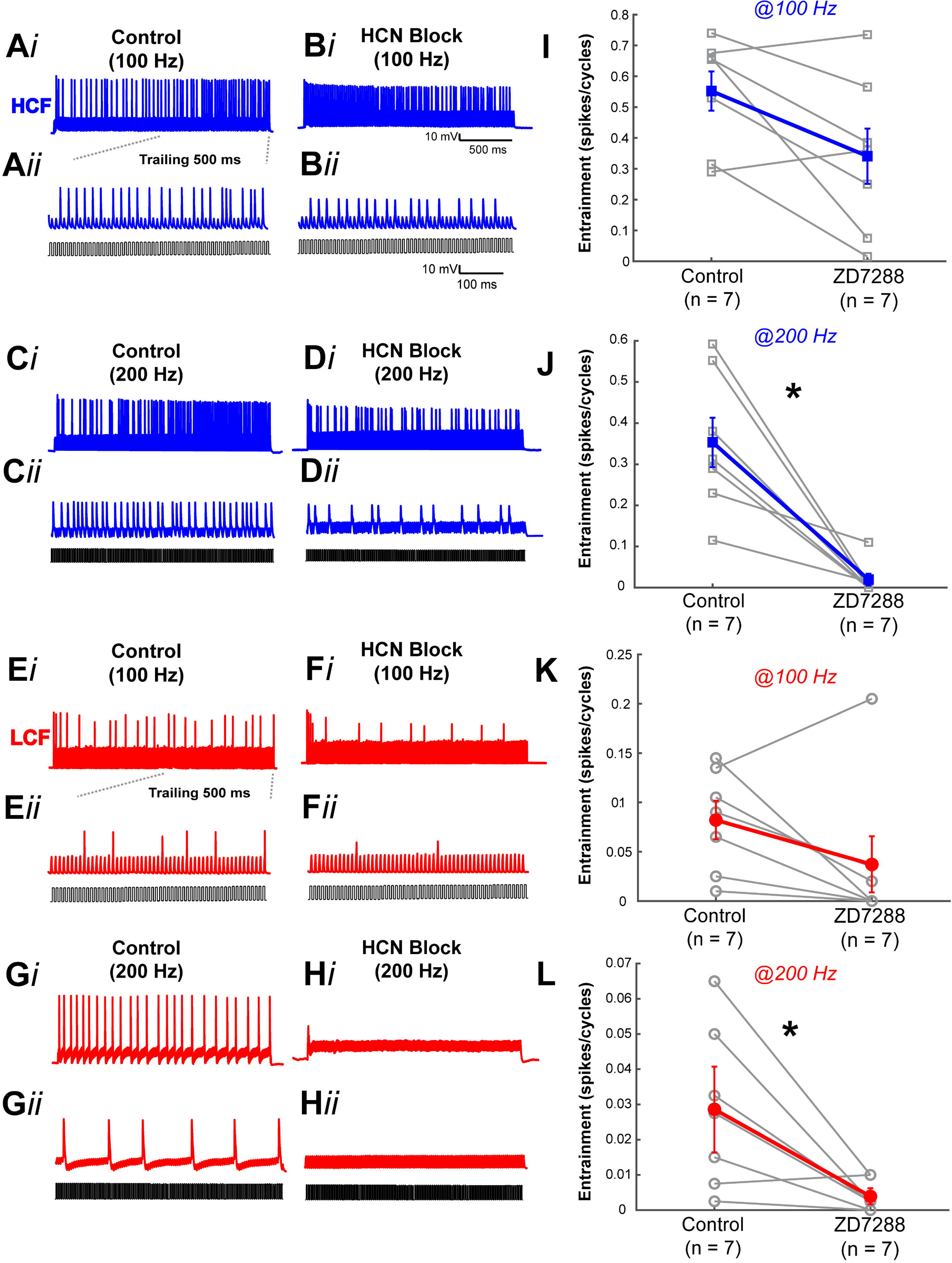
HCN channels enable action potential entrainment to depolarizing pulse trains in NM neurons. A*i*) Discharges of an HCF NM neuron to a 2s, 100 Hz pulse train in control and *Bi*) ZD7288 treatment. A*ii,* B*ii*) show the trailing 500 ms of the sustained portion of the response under each condition, which exhibit decreased firing rates during drug treatment. C&D) Voltage response of an HCF NM neuron to a 200 Hz square pulse train in control *(Ci)*, and ZD7288 treatment (*Di)*. C*ii* & D*ii)* show the trailing 500ms of the discharge responses under each condition. E&F) Response of an LCF NM neuron to a 2s, square pulse train at 100Hz in control (*Ei*) and ZD7288 treatment (*Fi*). The trailing 500ms of the discharge responses under each condition are zoomed in to show the sustained response (*Eii, Fii).* G&H) Voltage response of an LCF NM neuron to a 200Hz square pulse train in control (*Gi*) and ZD7288 treatment (*Hi*). The The trailing 500ms of the voltage response under each condition is zoomed in to show the sustained response (*Gii, Hii*). I,J,K,&L show entrainment data for each population under each condition as indicated. There was a decline in entrainment responses at both input rates for HCF and LCF neurons, however only the 200 Hz reductions were significant (LCF: *p<0.05, HCF: **P<0.0004, student paired t-tests).

**Table 1:**
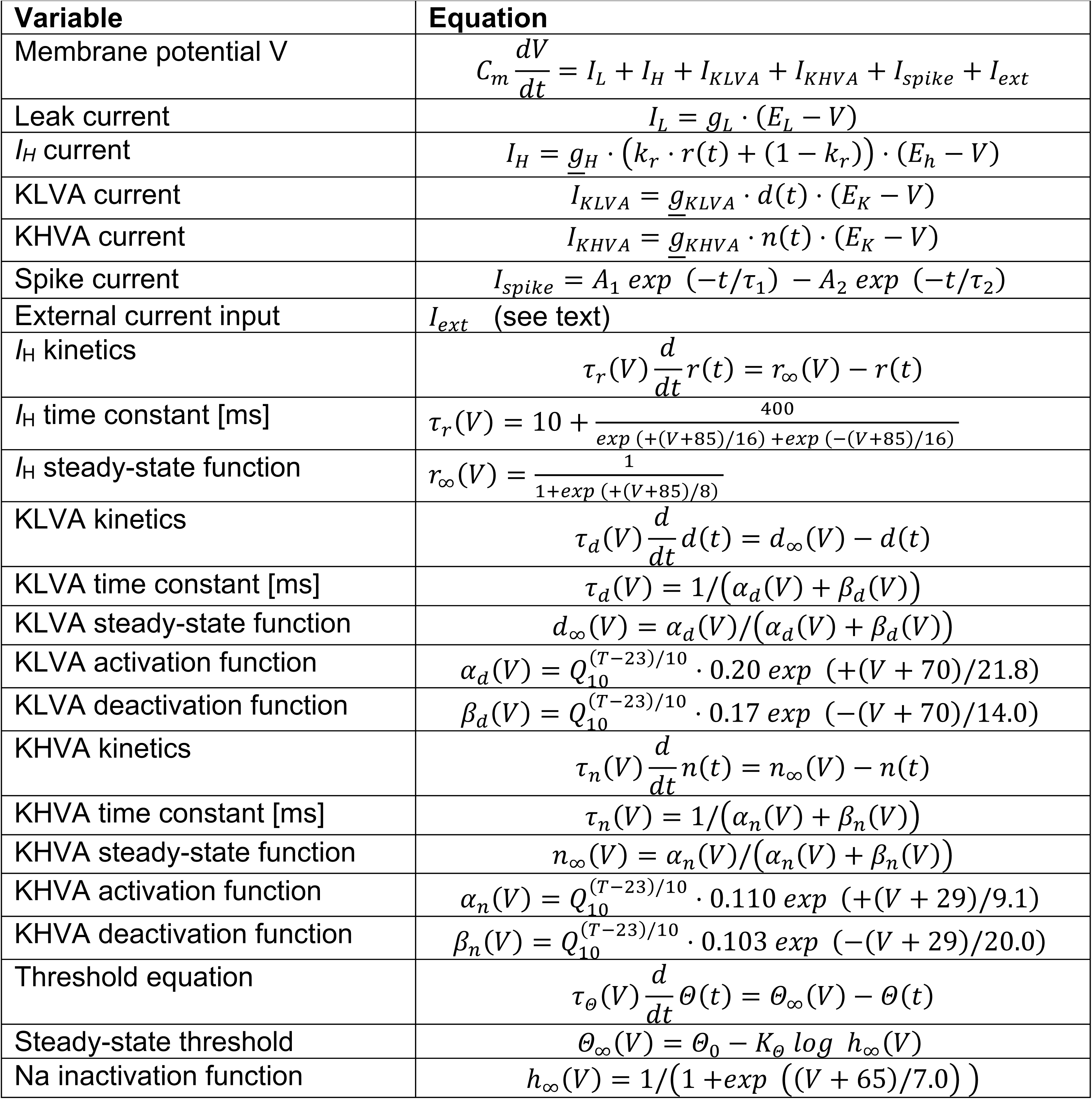
NM model equations. This model is based on the model used in Oline et al. (2016) and expanded with IH values derived from empirical data

**Table 2.**
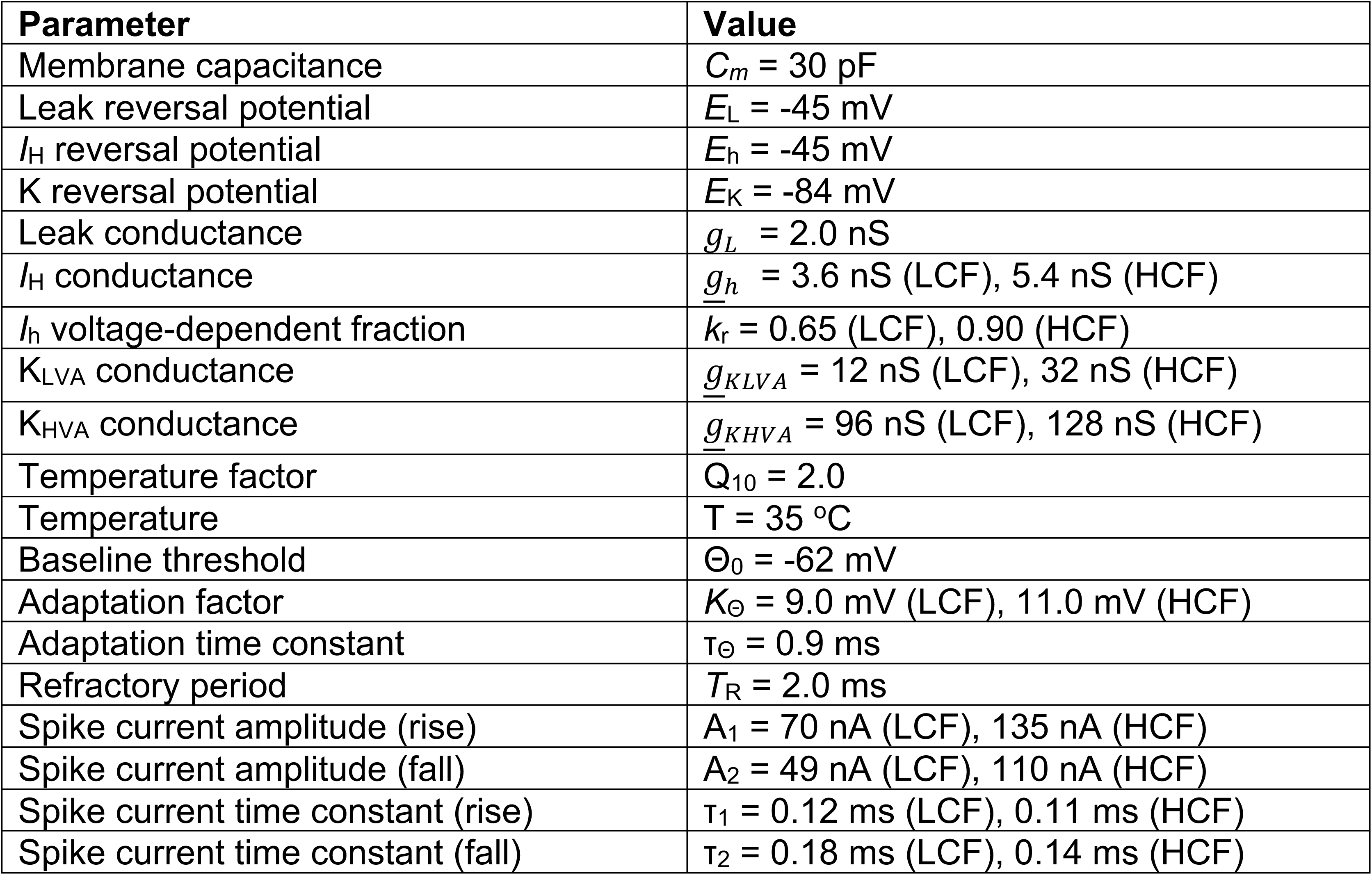
NM model parameters. . Note: entries with only one value indicate a common value for both LCF and HCF models

| Parameter | Value |
| --- | --- |
| Membrane capacitance | $C_m = 30$ pF |
| Leak reversal potential | $E_L = -45$ mV |
| $I_H$ reversal potential | $E_h = -45$ mV |
| K reversal potential | $E_K = -84$ mV |
| Leak conductance | $g_L = 2.0$ nS |
| $I_H$ conductance | $g_h = 3.6$ nS (LCF), 5.4 nS (HCF) |
| $I_h$ voltage-dependent fraction | $k_r = 0.65$ (LCF), 0.90 (HCF) |
| $K_{LVA}$ conductance | $\underline{g}_{KLVA} = 12$ nS (LCF), 32 nS (HCF) |
| $K_{HVA}$ conductance | $\underline{g}_{KHVA} = 96$ nS (LCF), 128 nS (HCF) |
| Temperature factor | $Q_{10} = 2.0$ |
| Temperature | $T = 35$ °C |
| Baseline threshold | $\Theta_0 = -62$ mV |
| Adaptation factor | $K_\Theta = 9.0$ mV (LCF), 11.0 mV (HCF) |
| Adaptation time constant | $\tau_\Theta = 0.9$ ms |
| Refractory period | $T_R = 2.0$ ms |
| Spike current amplitude (rise) | $A_1 = 70$ nA (LCF), 135 nA (HCF) |
| Spike current amplitude (fall) | $A_2 = 49$ nA (LCF), 110 nA (HCF) |
| Spike current time constant (rise) | $\tau_1 = 0.12$ ms (LCF), 0.11 ms (HCF) |
| Spike current time constant (fall) | $\tau_2 = 0.18$ ms (LCF), 0.14 ms (HCF) |

As in our experimental protocol for testing entrainment, we injected the same repeated current pulse trains (200 Hz, each 2-ms long rectangular pulse) to the model NM neuron (**Table 2**). The amplitude was fixed to 50 pA above threshold to simulate peri-threshold input conditions. To mimic membrane voltage fluctuations in vivo, we generated a wideband Gaussian noise and filtered it with a synaptic current filter of the form (−𝑡/𝜏_𝑅_) +𝑒𝑥𝑝 𝑒𝑥𝑝 (−𝑡/𝜏_𝐹_), with the rise and fall time constants being τR = 0.20 and τF = 0.33 ms, respectively (Oline and Burger, 2014). This filtered noise was then rescaled and injected as noise current to the model, so that the amplitude of membrane voltage fluctuation (measured by standard deviation) would match the desired value (0-2.5 mV; see Results). In our preliminary simulations, we also used low-pass-filtered Gaussian noise (cutoff frequency = 2 kHz), and found no qualitative differences from our model results.

## Results

We performed whole-cell voltage and current clamp experiments on 147 tonotopically categorized NM neurons from 39 late embryonic animals (aged E16-19). IHC analyses were performed on an additional cohort of 12 (P3) chicks. Computational modelling assessed the effects of the distribution of HCN channels on stimulus encoding capabilities of NM neurons across the tonotopic axis and under a broad range of input conditions.

### Prominent Voltage sags are a characteristic feature of the physiological response of NM neurons and mediated by HCN channels

We investigated differences among NM neurons’ membrane excitability across the tonotopic axis by injecting a range of hyperpolarizing and depolarizing 200 ms current steps (Range -200 to 500 pA, in 50 or 100 pA steps) in whole-cell current clamp (**Fig. 1**). We recorded the membrane voltage responses from 7 HCF and LCF NM neurons before and during application of the HCN antagonist, ZD7288 (40 µM). Sustained hyperpolarizing current steps evoked a slowly developing depolarizing sag particularly during the most hyperpolarized voltage responses for both a HCF and a LCF NM neurons (**Fig. 1 A, D)**. Depolarizing sags at hyperpolarized potentials usually indicate the onset of an inward current through hyperpolarization-activated cation channels (HCN). In order to test whether the depolarizing sags were due to hyperpolarization induced gating of HCN channels, we repeated the current step protocol in the presence of the HCN antagonist. During application, the depolarizing sag at hyperpolarized potentials was reduced or abolished in both HCF neurons (**Fig. 1 B, C**: HCF: n = 7; Control; sag = 38.3201 ± 3.14 %; ZD-7288(40 uM); sag = 6.9869 ± 2.89%; p = 1.91 x 10^-5^; paired t-test) And LCF neurons (**Fig. 1 E, F**; LCF: n = 7; Control; sag = 37.1575 ± 6.97 %; ZD-7288(40 uM); sag = 12.0909 ± 2.83 %; p = 0.0094; paired t-test)

### HCN conductances are tonotopically distributed in the NM

To further confirm the expression of HCN channels putatively underlying the voltage sags, we recorded membrane current responses to long duration (400 ms) voltage steps ranging from -20 to -120 mV in whole-cell voltage clamp in 10 mV increments (**Fig. 2**). The voltage protocol was designed to evoke responses across as much of the channels’ voltage activation range as possible, without disrupting membrane stability during the long-duration steps required to observe full activation. **Figure 2A and B** shows current responses from representative HCF and LCF neurons’ current responses, respectively. During larger hyperpolarizing steps ∼(−70 to −120 mV), slowly developing inward currents were observed in both neurons *black arrows*, putatively indicating HCN channel gating. To confirm the HCN channel dependence of these currents, we applied the selective HCN antagonist ZD-7288 (40 um) to all neurons. We observed that the slowly developing inward currents were eliminated at all voltages *open arrowheads* for both HCF (**Fig. 2C**) and LCF (**Fig. 2D**) neurons. **Figure 2E & F** show the current magnitude differences for pre- and post-antagonist single traces at - 120 mV for each neuron. In all voltage steps, the HCN-dependent currents of the HCF population were significantly greater than those of the LCF population (**Fig 2G**) (HCF: n = 8; LCF: n = 8; p = 0.0031; two-way ANOVA).

### Immunolabeling of HCN-1 channel subunits confirms tonotopic channel expression

To our knowledge, there have been no previous studies demonstrating HCN channel expression in the NM. Our physiological results suggest that there is a tonotopic distribution of IH in NM underlying the strong IH conductances that we observed in HCF neurons. IHC staining using antibodies targeting the HCN1 channel subunit confirmed a tonotopic gradient of HCN1 protein expression in NM neurons in post hatch (day P3) chicks (**Fig. 3**). Fluorescence intensity within individual neuron boarders were sampled and analyzed within high, middle, and low frequency regions of interest inscribed on images that transected the broad tonotopic representation of NM. Anti-HCN1 labeling was most robust in high frequency NM neurons (**Fig. 3E, H**) with lower expression in the middle and low frequency regions (**Fig. 3E-G), complementing voltage clamp recordings**. We quantified HCN1 expression by measuring pixel intensity within neuron boarders where intensity was normalized to the highest value observed in each tissue sample. Average pixel intensity within HCF NM regions was significantly higher than middle frequency (HCF: 84.57%, MCF: 56.19%, n = 4; p = 6.965 X 10 ^-5^, paired t-test) or low frequency (HCF: 84.57%, LCF: 45.03% p = 0.0004, paired t-test). Middle frequency region was also significantly higher than the low frequency region (p = 0.0413, paired t-test) (**Fig. 3C**). To confirm the specificity of the antibody, we performed pre-adsorption experiments using the antigen protein for HCN1. When the antibody was preincubated with the HCN1 protein antigen, staining was abolished (**Fig. 3I**) as confirmed by pixel intensity measurements (**Fig. 3D**). Tissue incubated with antibody alone had significantly higher pixel intensity values than tissue incubated in the presence of pre-adsorbed antibody (n = 4; p = 0.002064, paired t-test).

### HCN channels contribute to the tonotopic distribution of membrane excitability

To investigate the contribution of HCN channels to excitability, we injected depolarizing current ramps to NM neurons and measured action potential halfwidth, slope threshold, and integration period before and during ZD-7288 application at rest (**Fig. 4A**) (see Methods). Previous work demonstrated that slope threshold and integration period values recorded in response to systematically varying ramp stimuli provide experimentally efficient and robust measures of membrane excitability (McGinley and Oertel, 2006; Oline et al., 2016). Action potential halfwidth significantly increased about twofold during ZD-7288 treatment for HCF neurons (Control: n = 7, 0.26 ± 0.02 ms; ZD-7288: 0.53 ± 0.08 ms (40 uM), p = 0.0122, paired t-test) but a smaller magnitude increase observed in LCF neurons was not significant (control: n = 9; 0.52 ± 0.06 ms; ZD-7288: 0.63 ± 0.08 ms, p = 0.2269, paired t-test) (**Fig. 4C**). We also noticed a significant increase in integration period for HCF NM neurons (control; n = 7; 2.17 ± 0.13 ms, ZD-7288 2.87 ± 0.022 ms, p = 0.02; paired t-test) and not for LCF NM neurons (control; n = 7; 4.17 ± 0.47 ms, ZD-7288; 4.37 ± 0.60 ms in ZD-7288 ; p = 0.8839; paired t-test) (**Fig. 4D**). ZD-7288 application had no effect on the slope threshold measurements for both HCF (control; n = 9; 11.15 ± 1.12 mV/ms, ZD-7288; 10.09 ± 0.78 mv/ms; p = 0.4787; paired t-test) and LCF NM neurons (control, 6.38 ± 0.73 mV/ms, ZD-7288; 7.23 ± 1.02 mV/ms; p = 0.5622; paired t-test) (**Fig. 4B**).

However, we did not see an effect on action potential halfwidth, slope threshold or integration period during ZD-7288 application when these NM neurons were held at a voltage of -75mV.

### HCN channels enable NM neurons to reliably follow high synaptic inputs

Next, we asked whether HCN channels contribute to NM neurons’ encoding of temporal information. To address this question, we injected trains of depolarizing square-current pulses to NM cells at either a low (100 Hz) or high frequency (200 Hz) for 2 s epochs in control aCSF and in the presence of HCN antagonist. To ensure that current injection magnitude was appropriately scaled to the systematically varying excitability across NM, we first measured rheobase current in each neuron. Rheobase, or the current magnitude necessary to elicit a spike response in >50% of stimuli in increments of 50 pA to 100 pA, differed for HCF and LCF populations (LCF: 178.6+/-36.4 pA, HCF: 385.7+/-50.2 pA, p = 0.0107; paired t-test), consistent with expectations. We recorded the action potential responses to 100 Hz pulse trains, where pulse amplitudes were set at the empirically determined rheobase values for each neuron. **Figure 5** shows voltage traces to a 2s 100 Hz train in representative HCF and LCF neurons (**Fig. 5 *Ai, Ei*)**.

Typically, neurons responded with reliable onset responses followed by vigorous sustained spiking. In order to assess the neurons’ capacity to respond to long-duration periodic stimuli, we measured entrainment, the number of spikes produced per cycle. We excluded the first 100 ms of the response train from analysis as the onset responses were not affected. During application of 40 µM ZD-7288, onset responses persisted, but the sustained portion of the response exhibited moderately reduced spiking to low frequency input in both populations (**Fig. 5 Bi, Fi)**. Population data trended toward reduced entrainment for HCF neurons (HCF: control: 0.55 ± 0.06 spikes/cycle; ZD-7288: 0.34 ± 0.09 spikes/cycle, p = 0.85; n = 7; paired t-test) (**Fig. 5I**). Results were similar for LCF neurons despite lower response rates overall (LCF: control: 0.08 ± 0.02 spikes/cycle; ZD-7288: 0.04 ± 0.04 spikes/cycle, p = 0.38; n = 7; paired t-test) (**Fig. 5K**). While these reductions were not statistically significant, significant response suppression was observed at 200 Hz for both HCF neurons (**Fig. 5J**) (HCF: control: 0.35 ± 0.06 spikes/cycle; ZD-7288: 0.02 ± 0.01 spikes/cycle, p = 0.0003; n = 7; paired t-test), and LCF neurons (**Fig. 5L**) (LCF: control: 0.03 ± 0.01 spikes/cycle; ZD-7288: 0.004 ± 0.002 spikes/cycle, p = 0.02; n = 7; paired t-test).

### The addition of *I*H improves entrainment in a single compartment NM model neuron

In order to extend our physiological findings to investigate the IH contributions to entrainment in the presence of probabilistic input such as noise, we applied repeated pulse currents to a computational NM model neuron as in our experimental protocol. To mimic the fluctuation of the membrane potential *in vivo*, a synaptic-filtered Gaussian noise current of varied amplitudes was added to the stimulus pulse train (see Methods). The amplitude of the noise was measured by the standard deviation (SD) of the fluctuation in the membrane potential. For 100-Hz stimulation, the HCF model fired at almost every pulse cycle (**Fig. 6A**). This response pattern did not change even when all *IH* current was eliminated (**Fig. 6B**). The degree of entrainment, measured as the number of spikes divided by the number of pulses in 2000-ms stimulation, gradually decreased with the amount of noise (**Fig. 6E**). For noise SD over 1 mV, the HCF models showed a marginally lower entrainment than the LCF models, likely because of the stronger KLVA current and threshold adaptation, which generally hinder repeated spikes. For 200-Hz stimulation, the HCF model could not fire at every pulse cycle (**Fig. 6C**). The failure rate depended on the noise amplitude (**Fig. 6F**, filled circles). With all *IH* current removed (+ZD condition), the HCF model showed an increased failure rate (**Fig. 6D**). Removal of *IH* affected the entrainment for a wide range of noise level (**Fig. 6F**, open circles). This effect was also observed for the LCF model (**Fig. 6G**). For large noise levels, the difference between the control and ZD condition became small, because the spiking behavior of the models was dominated by the noise rather than the input pulses. These modeling results suggest that *IH* may help NM neurons entrain to high-frequency, peri-threshold stimulation.

**Figure 6.**
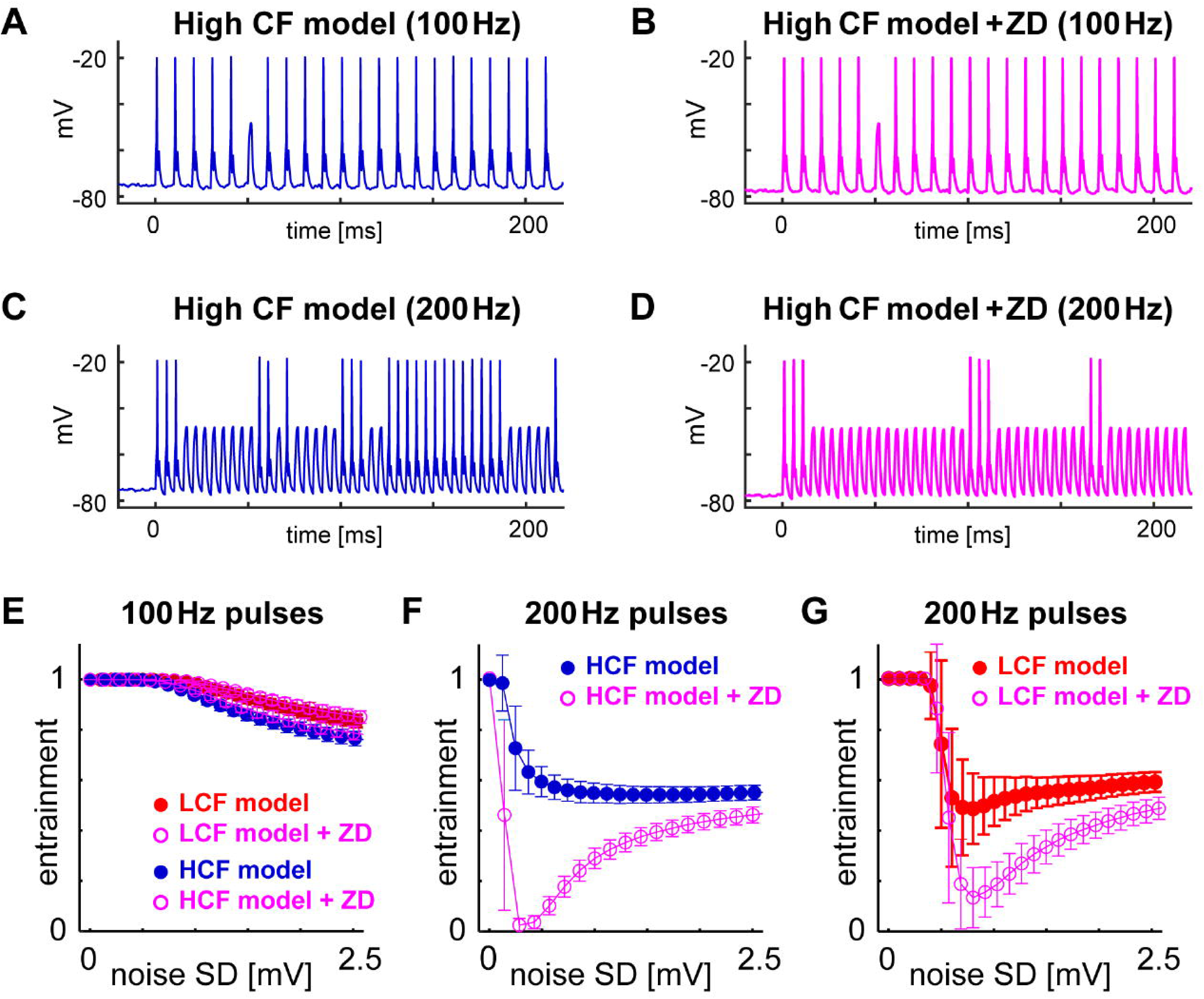
Model high CF neuron shows a reduction in entrainment when IH is removed. Sample voltage trace for model HCF neuron at (*A&B*) 100 Hz and (*C&D*) 200 Hz before and after removing the IH conductance. E) NM model neurons are able to entrain to 100 Hz inputs regardless of noise fluctuations with and without IH. F&G) Entrainment is reduced for both the low and high CF model neurons when the IH conductance is removed.

## Discussion

In this report, we demonstrated that HCN1 channels are tonotopically expressed within the avian cochlear nucleus, and are most abundant in HCF neurons. We determined the relative contribution of *I*H to membrane excitability using depolarizing ramp input stimuli from potentials near rest and found that blocking *I*H increased Action Potential halfwidth and integration periods for HCF neurons. Finally, we tested the potential functional contribution of *I*H using sustained pulse train stimuli as a proxy for sound driven periodic synaptic input both in model and actual neurons. Both experimental and modeling data suggest that *I*H regulates action potential entrainment for HCF neurons and thus contribute to response fidelity specifically during high-frequency stimuli. These results suggest that *I*H contributes to excitability optimization that enables reliable response fidelity to high frequency patterned synaptic input. These data are the first comprehensive analysis of *I*H distribution and function across the tonotopic axis of NM.

HCN channels are expressed throughout the central and peripheral nervous systems. However, their function within a particular region is often challenging to interpret because they can contribute both to enhancement or suppression of activity (Accili et al., 2002; Kase and Imoto, 2012; Ventura and Kalluri, 2019). One feature of HCN channels that contributes to this functional diversity is the ligand-dependent modulation of this voltage-gated channel. HCN channels include a cAMP sensing domain, which when bound shifts HCN activation voltages to more positive potentials (He et al., 2014). Of the four HCN channel isoforms, HCN1 channels are less responsive to cAMP binding (He et al., 2014). Within NL cAMP narrows the coincidence detection window specifically for HCF neurons, which suggests that *I*H may be modulated as a means to improve coincidence detection (Yamada et al., 2005).

HCN conductances are partially activated near resting voltages, thus providing a steady inward depolarizing current that sustains membrane voltage closer to action potential threshold, in opposition to a predominant low voltage activated outward potassium conductance. Crucially this partial activation also decreases membrane resistance and thus opposes membrane voltage fluctuations. High density expression of IH channels has been shown to generate substantial membrane voltage noise which significantly impacts action potential fidelity in modeled pyramidal neurons (Kole et al., 2006). Incorporating membrane voltage noise within our NM model, showed that *I*H is especially vital for ensuring action potential fidelity during moderate stochastic fluctuations.

In the mammalian medial superior olive (MSO), *I*H and low voltage-activated potassium channels cooperatively regulate membrane excitability to maintain consistent responses to simulated synaptic input (Khurana et al., 2011; Baumann et al., 2013). In the current study, we examined the relative influence of HCN on threshold stability when challenged with time-varying ramp input. Blocking HCN channels increases integration periods for high CF neurons. These observations are consistent with observations from McGinley and Oertel (2006), which revealed a negative correlation between the magnitude of the *I*H related voltage sag in response to hyperpolarizing current and the integration window. These findings support the assessment of the relative interactions between *I*H and KLVA as they relate to controlling the temporal encoding abilities of NM neurons.

Furthermore, our in-vitro and modeling results indicate that *I*H enhances sustained entrainment to periodic inputs during long-duration stimuli. These data complement the growing body of work in several sensory processing regions that also determine that *I*H is essential for maintaining spike fidelity in various areas of the nervous system (Kole et al., 2006; Endo et al., 2008;Baginskas et al., 2009; Byczkowicz et al., 2019). In short, blocking HCN channels in vestibular ganglion neurons (VGN) significantly reduces spontaneous firing (Horwitz et al., 2014). However, others show that enhanced HCN channel activation by cAMP increases spike count but reduces spike regularity in VGNs (Ventura and Kalluri, 2019). Thus, future studies may investigate whether further activation of *I*H enhances or dampens potential action entrainment within NM.

In conclusion, HCN channels and *I*H currents are tonotopically distributed with NM in a pattern that mimics the expression of other voltage gated channels in this system (Fukui et al., 2004; Burger et al., 2011; Adachi et al., 2019). Functionally, this distribution appears to enable spike regularity of HCF NM neurons to periodic input especially in the presence of noise. Taken together these data suggest that *IH* may be among several voltage gated channels that are highly regulated during development to optimize temporal encoding.

## Acknowledgements

We would like to thank Drs. Julie Haas and Paul Asadourian. This work was supported by NIH/NIDCD Grants DC-008989 and DC 018537. GA’s work is supported by the Deutsche Forschungsgemeinschaft (DFG: EXC2177/1, Project ID 390895286) and an Associate Junior Fellowship by the Hanse-Wissenschaftskolleg.

## Supplementary

**Supplementary Figure 1. Responses of NM neuron models to whole cell clamp protocols reproduced experimental data** A) Current Response of a LCF model (*top*), and a HCF model (*middle*) to simulated holding voltage steps (−20 to -120mV, in 10mV increments, *bottom*). B) I-V plot showing mean IH sensitive current responses for both LCF and HCF models. C) Representative voltage responses for a LCF model (*top*) to simulated ramp current injections (*bottom*). D) Representative voltage responses for a HCF model (*top*) to simulated ramp current injections (*bottom*). E) Slope threshold before and after removing IH conductance at rest, -80 mV, and -75 mV for LCF and HCF models. F) Integration period before and after removing IH conductance at rest, -80mV, and -75mV for LCF and HCF models.

