## Supplementary figures and images for "Hyperpolarization-activated cation channels confer tonotopic specialization for temporal encoding of sound frequency in the cochlear nucleus"

### Supplementary Figure 1

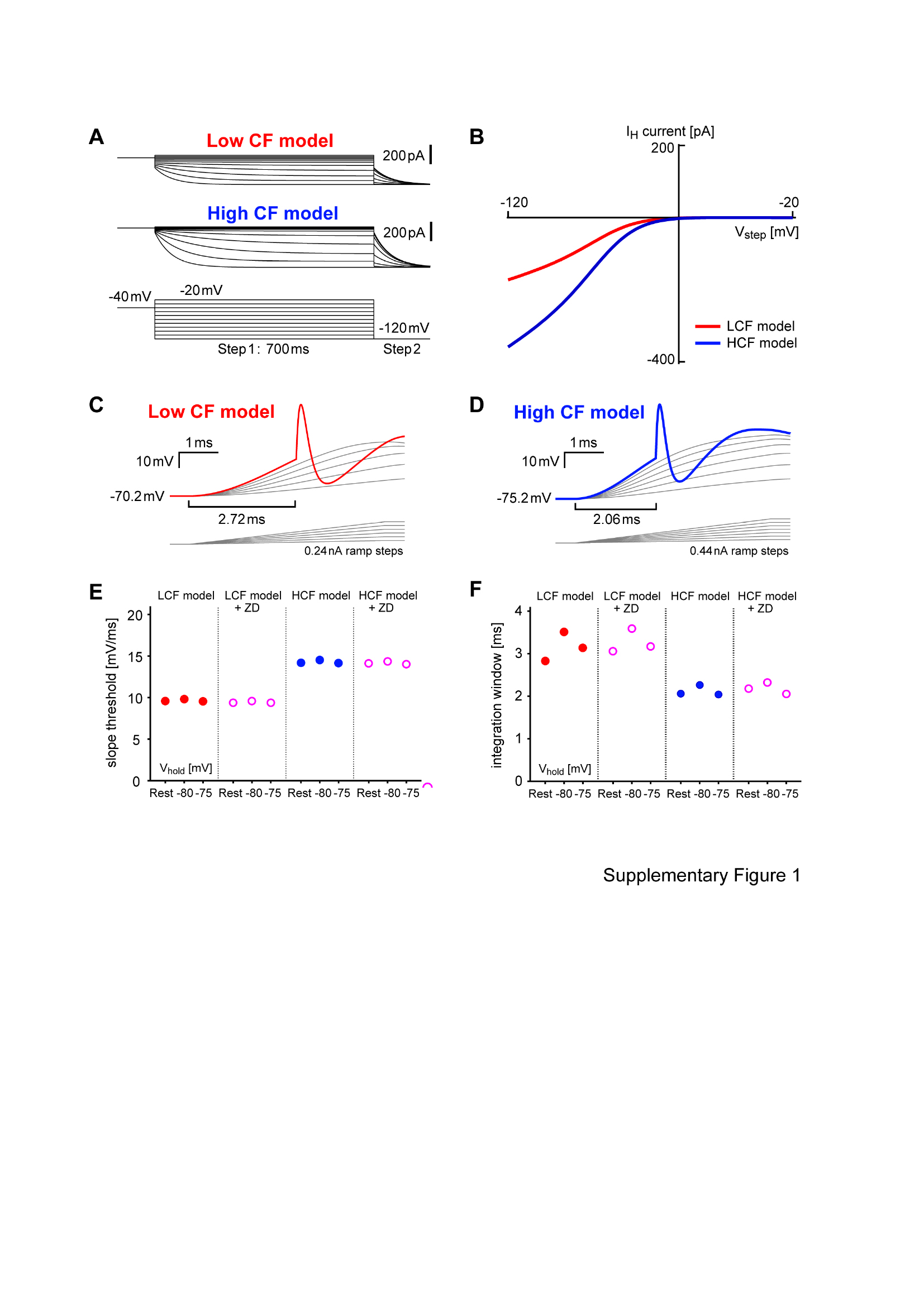
